# Maternal Care Leads to the Evolution of Long, Slow Lives

**DOI:** 10.1101/2024.01.05.574353

**Authors:** Matthew N Zipple, H Kern Reeve, Jimmy H Peniston

## Abstract

Animals, and mammals in particular, vary widely in their “pace of life,” with some species living long lives and reproducing infrequently (slow life histories) and others living short lives and reproducing often (fast life histories). These species also vary in the importance of maternal care in offspring fitness: in some species, offspring are fully independent of their mothers following a brief period of nutritional input, while others display a long period of continued dependence on mothers well after nutritional dependence. Here we hypothesize that these two axes of variation are causally related to each other, such that extended dependence of offspring on maternal presence leads to the evolution of longer lives at the expense of reproduction. We use a combination of deterministic modeling and stochastic agent-based modeling to explore how empirically-observed links between maternal survival and offspring fitness are likely to shape the evolution of mortality and fertility. Each of our modelling approaches leads to the same conclusion: when maternal survival has strong impacts on the survival of offspring and grandoffspring, populations evolve longer lives with less frequent reproduction. Our results suggest the slow life histories of humans and other primates as well as other long-lived, highly social animals such as hyenas, whales, and elephants, are partially the result of the strong maternal care that these animals display. We have designed our models to be readily parameterized with demographic data that is routinely collected by long-term researchers, which will facilitate more thorough testing of our hypothesis.

**Significance Statement:** Humans and other primates live longer lives and reproduce less often than other mammals of similar body mass. What is the cause of these long lives? Here we add to existing hypotheses, including the Mother and Grandmother hypotheses, by arguing that these increased lifespans are partially explained by the intense maternal care that many primates express. Using a combination of deterministic and stochastic modeling approaches, informed by empirical data, we show that stronger connections between maternal survival and offspring fitness leads to selection for longer lives and slower reproduction. Our models suggest that the importance of the mother-offspring relationship, which defines much of human and non-human primate lives, lies at the core of the evolution of our long lives.

## Introduction

The dependent relationship between offspring and mother is a defining feature of mammalian life history (1). The nature and timing of this dependent relationship can vary substantially between species (2, 3). For example, species with rapid generation times (such as mice and many other rodents) are generally assumed to achieve complete independence from their mothers at the termination of milk transfer, such that dependency terminates at weaning (3–5). In contrast, in other mammalian species mothers continue to provide critical social, nutritional, or informational inputs for their offspring (e.g. primates (2, 6–8); cetaceans (9); hyenas (10); elephants (11)). The result is that maternal presence or absence in an offspring’s life up to (and sometimes past) the age of sexual maturity has a major impact on offspring fitness in these species. In such species, a mother’s survival at any given age is intertwined with the fitness of not only those recently-born offspring that depend on their mother completely for milk, but also those previous offspring who are no longer nursing but still rely on their mother in other ways and who might be several years old (or even a decade or more in bonobos, (12, 13)).

Mammals also vary markedly in ‘pace of life,’ a phenotype that captures correlated variation in life history traits that can range from ‘slow’ to ‘fast’ life histories (14). Slow life histories are characterized by a late age at maturity, infrequent reproduction, and long lifespans, while species with fast life histories mature early, reproduce often, and die young (15). These traits necessarily co-vary due to tradeoffs between investment in survival and reproduction (16–18). Pace of life is strongly predicted by body size (14, 18, 19), but a great deal of variation exists around this trend. And primates, as a group, live strikingly longer, slower lives than non-primates of comparable body size (14).

How should variation in offspring dependence on mothers shape the evolution of mammalian pace of life? When maternal longevity influences offspring survival, a mother’s investment in its own survival has an indirect genetic effect on offspring’s fitness (20, 21). Because of the relatedness of mother and offspring, we expect this social selection to lead to increased positive selection on maternal survival (20, 21). This logic has been previously captured in the grandmother and mother hypotheses (22–25) that seek to explain extended post-reproductive lifespans in those species in which they occur (i.e. humans and whales; (9, 26, 27)). According to these hypotheses, survival even in the absence of reproduction enhances maternal fitness because mothers are able to increase the fitness of their offspring and grandoffspring. Yet, these hypotheses focus on the most extreme outcomes observed— long lives featuring an extended period of complete infertility in later life (menopause) that characterize only a very narrow range of mammalian species (even among primates, humans alone display menopause (26)).

Here we seek to describe a more general form of this logic, which should apply across the mammalian taxonomy: as offspring fitness becomes more tightly linked to maternal survival, or as this linkage becomes extended across wider ranges of an offspring’s life, mothers should experience stronger selection to survive, so as to promote offspring fitness. Given constraints on mothers’ resources, this selection should lead to an inevitable tradeoff, such that mothers live longer at the expense of a lower reproductive rate (14, 18). In other words, we hypothesize that extended maternal care should lead to the evolution of the long, slow life histories that characterize primates and several other highly social orders of mammals.

The purpose of this paper is to understand how this extended dependence (hereafter “mother-offspring fitness links”) shape the evolution of mammalian life history. Here we (1) formalize the above qualitative model in a general form, (2) build a model of mammalian life history that specifically incorporates observed mother-offspring fitness links, and (3) use an evolutionary agent-based model to test predictions about how mother-offspring fitness links should lead to the evolution of long lives at the expense of reproduction.

### Empirical Links Between Maternal Survival and Offspring Fitness

In a recent study of seven non-human primates, Zipple and colleagues describe four testable predictions about the ways in which maternal death (or survival) during an offspring’s immature period should shape that offspring’s fitness throughout its life (28). These predictions were designed to be testable with demographic data that are routinely collected in dozens of long-term studies of free-living mammals (29, 30). We adopt this same approach here, with the aspiration that our model can be readily tested with similar demographic data. Because of their centrality to this paper, we now reproduce and expand on the description of the mother-offspring fitness links proposed by Zipple and colleagues (2021).

Imagine a young immature mammal (F1), whose mother (M) has just died. What impacts will that death have on F1’s fitness? First, if F1 is not yet at the age of weaning at the time of M’s death, F1 will almost certainly die. We define weaning as the age at which it is possible for an offspring to survive in the absence of any milk transfer, meaning that maternal death prior to this age necessarily means that F1 will die, except in those relatively unusual species in which non-mothers often provide milk to offspring (e.g. white-faced capuchins (31), house mice (32)). We also assume that the death of M during the immature period has both acute and chronic negative impacts on F1’s condition. As a result, even if F1 is past the age of weaning (but still immature) at the time that its mother dies, we expect it to be at an increased risk of death for the remainder of life, including the rest of its immature period (Figure 1, blue arrow) as well as in adulthood (Figure 1, red arrow).

**Figure 1.**
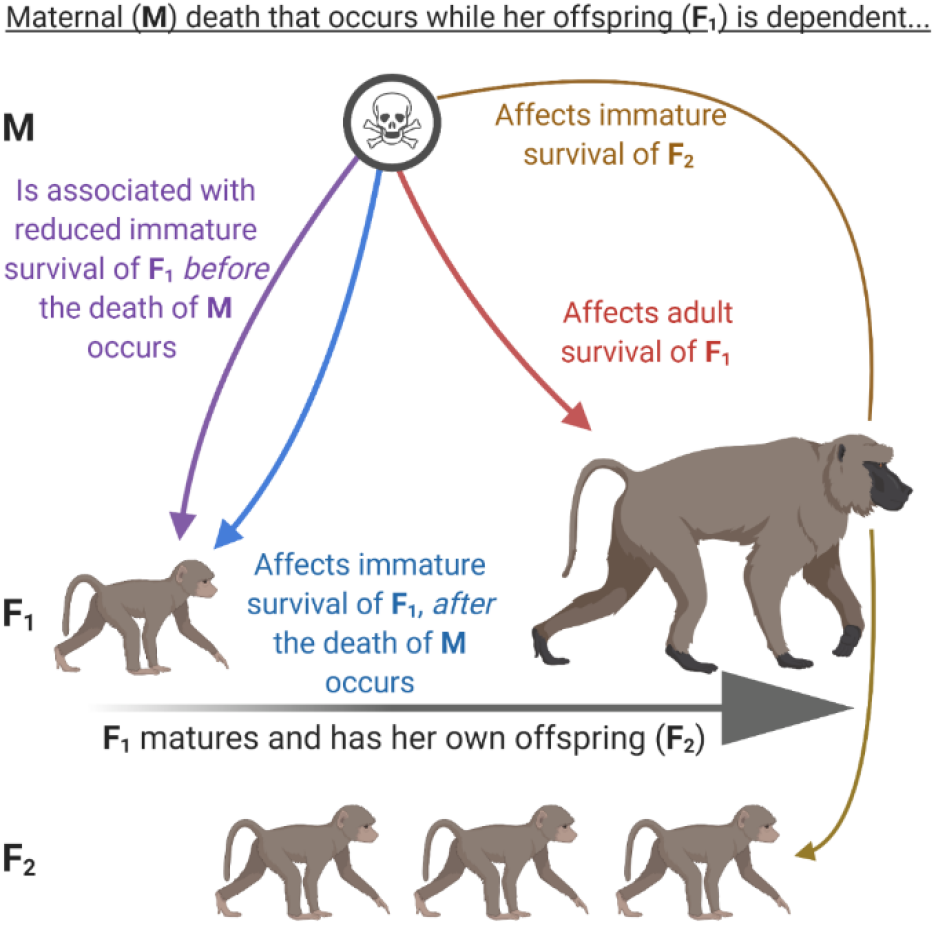
Four ways in which maternal death in an offspring’s early life is predicted to affect offspring fitness. Reproduced from Zipple et al (2021).

Finally, we assume that when an animal is in poor condition, it is more likely to die and less able to provide maternal care to its offspring. Thus, we predict that F1, who we assume to be in chronically worse condition in adulthood due to M’s death, will be less able to provide maternal care to its own offspring (F2). As a result, we predict an intergenerational effect on offspring survival of M’s death in F1’s early life (Figure 1, gold arrow). Finally, we assume that M’s death in F1’s early life was in part the result of M being in relatively poor condition during the years leading up to its death. We therefore predict that F1 will face an increased risk of mortality in the years preceding M’s death, when M is still alive (Figure 1, purple arrow).

Each of these predictions has been supported in the well-studied free-living Amboseli baboon and Gombe chimpanzee populations (28, 33, 34), and each individual prediction has received substantial support from a range of primate and non-primate mammals (8–11, 28, 35–38). Yet, even within great apes this pattern is not universal—gorilla offspring are remarkably robust to the loss of mothers, and some animals orphaned even just past the age of weaning do not appear to be any worse off as a result of losing their mother (28, 39). It is therefore clear that in some mammalian species, the survival prospects for a female and its offspring are dependent of the survival of its mother, while in others fitness outcomes of mother and offspring are much less interrelated (2, 3). The goal of the rest of this paper is to explore the implications of that realization for the evolution of pace of life histories.

We take three related approaches to considering how links between maternal survival and offspring fitness should shape the evolution of female pace of life. First, we present a general model of how maternal investment in survival and reproduction should be selected for, depending on the relationship between maternal survival and offspring survival. We then develop a more specific deterministic model, based on specific mammalian life history characteristics to show how this general model applies to life histories most similar to our own. Finally, we build a stochastic agent-based evolutionary model based on our deterministic model and parameterized based on empirical results from the Amboseli baboons to validate our predictions in an evolutionarily and ecologically relevant context.

### General Model

Let us assume a female’s fitness, *w* is a function of its survival, fertility, and the survival of its offspring. The female’s survival, *s*, is a function of its investment, *y*, in its own lifetime extension such that *s(y)* is a monotonically increasing function. The female’s investment in survival trades off with investment in fertility, *f*, such that *f(y)* is monotonically decreasing. Finally, we assume that offspring survival has some baseline likelihood, *a*, and that survival can be enhanced as a result of the mother surviving so that the mother can provide inputs for its offspring, thereby improving offspring survival chances. Thus, offspring survival is *a* + *so(y)*, where *so(y)* is a monotonically increasing function that captures the increase in offspring survival that maternal survival confers.

**Table 1.** Parameters used in our deterministic, mammalian life history model.

| Variable | Meaning |
| --- | --- |
| $w$ | Female fitness |
| $s(y)$ | Female survival as a monotonically increasing function of $y$, investment in survival |
| $f(y)$ | Female fertility as a monotonically decreasing function of $y$ |
| $a$ | Baseline offspring survival probability in the absence of maternal survival |
| $so(y)$ | The increase in offspring survival probability conferred by maternal survival, which is in turn a function of maternal investment in the mother's own survival. |

A female’s fitness can then be described by:

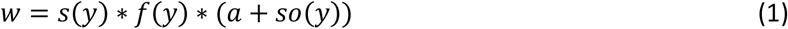

And the first derivative of fitness with respect to *y* is therefore:

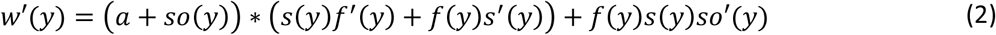

If we set the derivative to zero and solve for *f’(y)* at the maximum value of *w* we obtain:

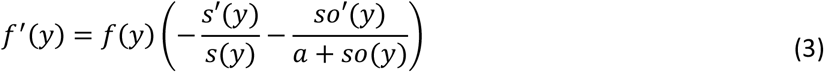

If we then take the derivative of *w’(y)* with respect to *a*, baseline offspring survival, we find:

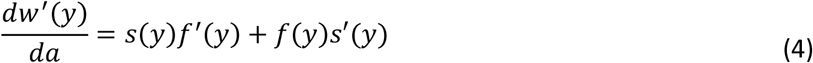

Substituting for *f’(y)* from above and simplifying we find:

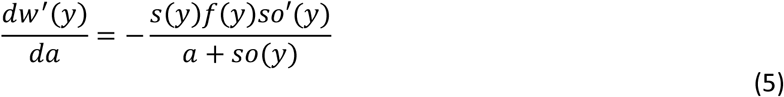

Because *so(y)* is always greater than zero and monotonically increasing, this derivative is negative. That is, as baseline survival of offspring increases, and the opportunity for mothers to improve offspring survival through their presence decreases, the optimal value of *y* declines. In contrast, when offspring survival in the absence of a mother decreases and maternal presence is able to substantially increase offspring survival, there is selection for increased maternal investment in the mother’s own survival, at the expense of reproduction.

### Mammalian Life History Model

Given the predictions from our general model, we next present a model that specifically considers the pathways by which mothers can improve offspring survival within the context of mammalian life history. This life history includes a period of offspring nutritional dependence on their mother until weaning age (*w*) and a juvenile phase between weaning at the age of sexual maturation (*b*). Individuals then reproduce with an age-related frequency until age *d*, when they either die or otherwise cease reproduction (e.g. at the onset of menopause).

**Table 2.**
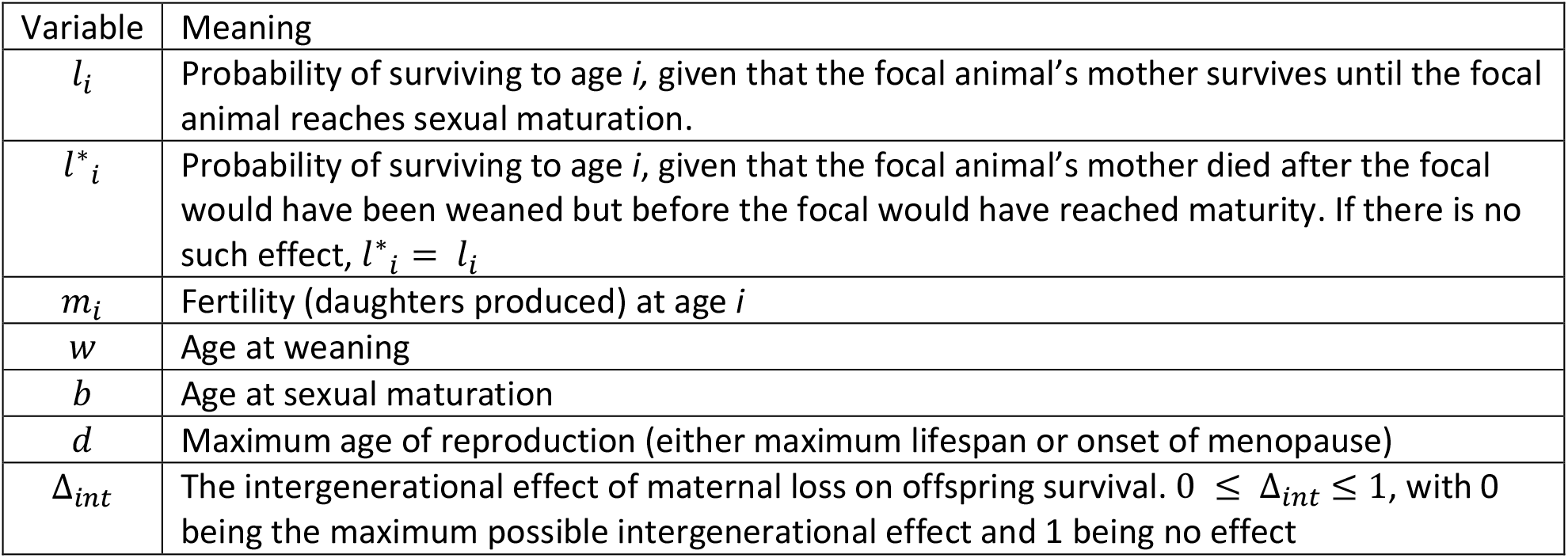
Parameters used in our deterministic, mammalian life history model.

**Table 3.** Scenarios considered in our mammalian life-history models.

| Scenario | Conditions |
| --- | --- |
| 1 | Offspring and grandoffspring survival are completely independent of maternal survival |
| 2 | Offspring die if their mothers die before offspring are weaned (age $w$) |
| 3 | In addition to (2), offspring are at an increased risk of death if they are born in the last $b$ years of their mothers lives. That is, if their mothers die before the offspring would become sexually mature. The magnitude of this impact may differ across juvenile (blue and purple arrows, Figure 1) and adult (red arrow, Figure 1) phases |
| 4 | In addition to (3), grandoffspring are at an increased risk of death if they are born to individuals who lost their mother prior to maturation. |

Within this mammalian life history context, we explore 4 different scenarios in which links between maternal survival and offspring and grandoffspring survival may exist to varying extents: (1) maternal survival and offspring survival are completely independent, (2) maternal death prior to offspring weaning results in offspring death, (3) in addition to the previous, maternal death after offspring weaning but before offspring sexual maturation leads to reduced survival of offspring in the juvenile period and/or adulthood, and (4) in addition to the previous, maternal death before offspring sexual maturation leads to an intergenerational effect on grandoffspring survival. Scenario 4 most closely matches results observed in several primates (28).

Because Scenario 4 involves a link between maternal survival and grandoffspring survival, we take the expected number surviving granddaughters that a female will produce as our proxy for fitness. Under Scenario 1, when maternal survival and offspring survival are independent, the number of granddaughters that survive to maturation that a female is expected to produce is:

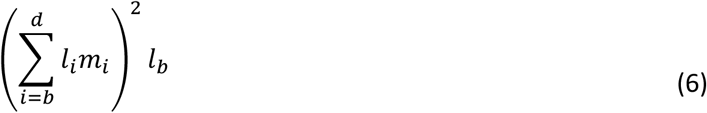

This equation is simply the number of daughters that a female is expected to produce, multiplied by the number of daughters that each of those daughters will produce and the probability of those offspring (granddaughters) surviving to maturity (age *b*).

Under Scenario 2, when maternal death prior to offspring weaning leads to offspring death, this equation is only slightly changed:

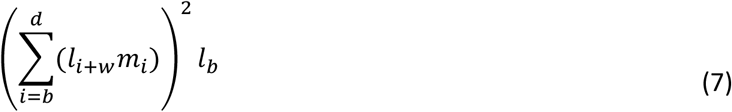

Here, all offspring born in the last *w* years of the a mother’s life perish. But if their mothers survive until these offspring are weaned, there is no longer any further link between their survival and the survival of their mothers.

Under Scenario 3, where maternal death that occurs between offspring weaning and offspring maturation has a permanent and ongoing negative impact on offspring survival, the situation becomes substantially more complicated, with the expected number of surviving granddaughters being:

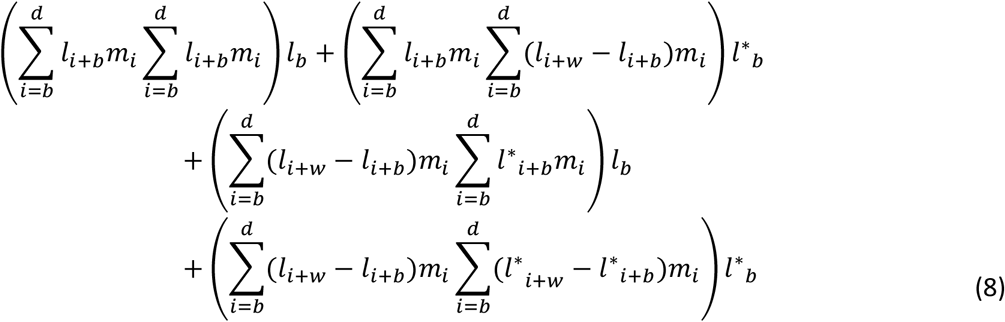

Here, daughters and granddaughters that are born more than *b* years before maternal death are unaffected by the mother’s later death (hereafter ‘uncompromised’ offspring). But any offspring born in the last *b* years of the mother’s life are at perpetually increased risk of death, denoted by *l*^*\**^ (hereafter ‘compromised’ offspring). Though *l*^*\**^ is always lower than *l*, the magnitude of the difference between the two might vary across the lifespan. Maternal loss might have a particularly strong effect during the remainder of the juvenile phase as compared to adult survival, for example. The first two terms in equation (8) correspond to the survival of granddaughters that are born to uncompromised daughters and the second two terms correspond to survival of granddaughters born to compromised daughters. These granddaughters may themselves be compromised (the second and fourth terms) by the death of their own mothers.

Finally, under Scenario 4, granddaughters born to compromised daughters face an additional intergenerational effect of maternal loss, regardless of when their mothers (the ‘compromised daughters’) actually die. Here the number of surviving granddaughters that an animal produces is very similar to Scenario 3, with the addition of an intergenerational modifier, Δ_*int*_ being applied to all granddaughters born to compromised daughters.

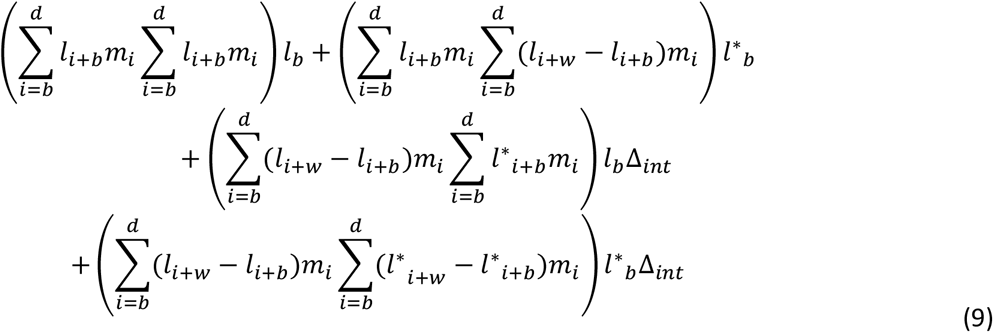

In each step from Scenario 1 through Scenario 4, the ultimate cost of death to an animal’s inclusive fitness increases because of the downstream impacts that death will have on its daughters or granddaughters. If we assume the presence of a tradeoff between an individual’s investment in its survival and its investment in reproduction—as measured through fertility—we would predict that females living under Scenario 4 should be selected to increase investment in survival at the expense of reproduction. Note that this is the same qualitative conclusion reached by our general model.

We next quantify this prediction by parameterizing two models that we built using published life table data from the Amboseli baboon population (29). We chose this population as our example because of the large sample size of animals whose lives’ data generate the life table and because it is the system in which the links between maternal survival and offspring and grandoffspring survival are best documented. We emphasize, however, that the results of our model should generalize to any system where such mother-offspring fitness links exist.

Our first model is deterministic and proceeds by three steps. First, using the above equations, we calculated the number of surviving granddaughters that a female would be expected to produce under each of Scenarios 1-4, using published life history data (hereafter *g*_*0*_). We then perturbed the birth rate of this system by either increasing or decreasing annual fertility from its baseline (by multiplying by annual fertility by a constant, *δ*_*m*_) and calculated the number of surviving granddaughters that a female would be expected to produce under each scenario given this new birth rate (hereafter *g*_*1*_). Finally, we calculated the amount that adult mortality would need to increase or decrease alongside birth rate under each scenario (by multiplying adult mortality by a constant, *δ*_*q*_), such that *g*_*1*_ = *g*_*0*_. This value (hereafter *λ*) represents the magnitude of the tolerable tradeoff between investment in survival and reproduction that could be selected for.

Our final model is a stochastic agent-based evolutionary model that is meant to assess the predictions of the deterministic model in an evolutionarily relevant population. A full description of this agent-based model is available in the supplement. Briefly, we simulated populations of females that followed the same published baboon life history table used in the deterministic model. Females proceed through life in one-year time steps, probabilistically dying and reproducing. After a 200-year burn-in period, we introduced a mutation to 50% of the population that changed both birth rate and death rates by multiplying them by *δ*_*m*_ and *δ*_*q*_ respectively. We allowed the populations to evolve for an additional 1000 years or until the mutation was either fixed or lost. For each value of *δ*_*m*_, we calculated (via linear interpolation) the value of *δ*_*q*_ that corresponded to 50% of the populations becoming fixed with the mutation (i.e. the value at which selection was neutral). This value is equivalent to *λ* in the deterministic model.

Our deterministic and agent-based models yielded very similar results (Figure 2). When we increased birth rate from baseline, the magnitude of the associated mortality cost for which the mutation remained advantageous was lower when links between maternal survival and offspring survival were present (Scenario 4) as compared to when they were absent (Scenario 1). Similarly, when a mutation decreased birth rate from baseline, the magnitude of the associated reduction in mortality that was necessary for the mutation to be advantageous was lower under Scenario 4 as compared to Scenario 1. That is, the cost of a lower birth rate could be offset by a smaller reduction in adult mortality when mothers’ own survival shaped the survival of their offspring and grandoffspring. Thus, when links between maternal survival and offspring fitness were present, our simulated populations evolved longer lives and slower reproduction as compared to when they were absent.

**Figure 2.**
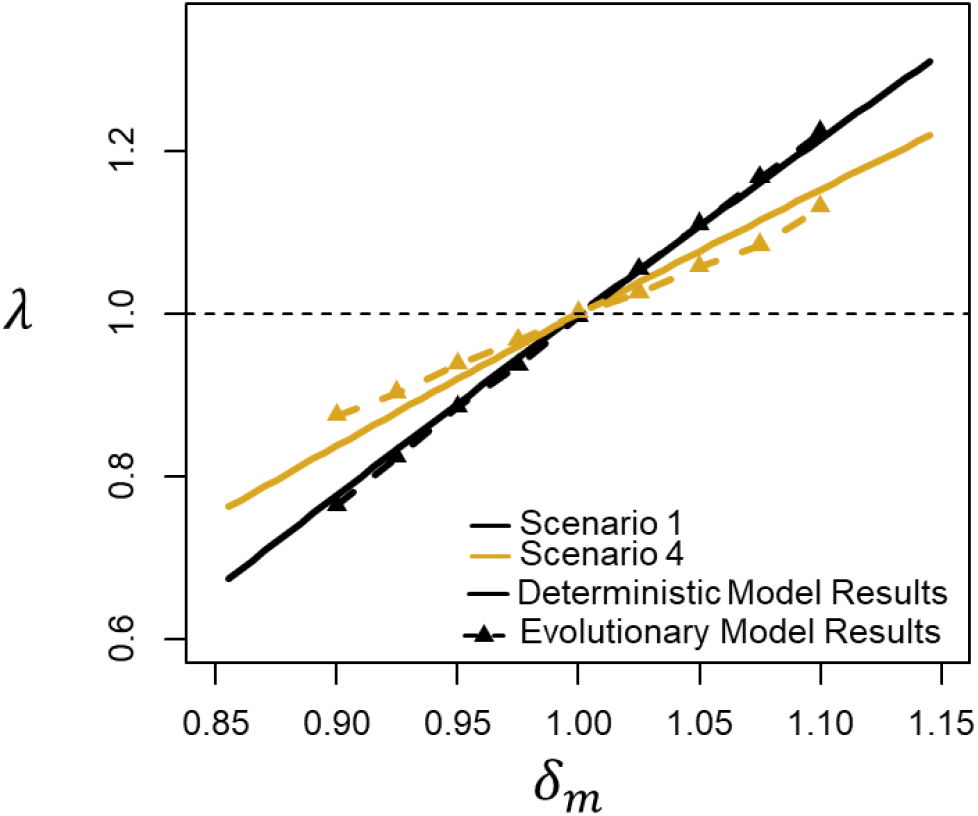
Deterministic and evolutionary, agent-based models of a population of baboon-like animals yield the same qualitative result. Specifically, for a given change in birth rate, the change in mortality rate necessary to result in neutral selection is smaller in magnitude when links between maternal survival and offspring fitness are present (gold, Scenario 4) than when they are absent (black, Scenario 1).

There was some quantitative discrepency between our deterministic and agent-based modeling results, with the values of *λ* that emerge from the stochastic agent-based model being constently more extreme than those that emerge from the determinstic model. This difference likely results from the agent-based model allowing for a more extended intergenerational cascade that we were not able to incorporate in our two-generation deterministic model. That is, in equation 9 we follow fitness outcomes only through the grandoffspring generation. But in reality, there will continue to be effects of a female’s premature death even beyond this generation, as each of her offspring are now more likely themselves to imperil the eventual survival of their grandoffspring (i.e. the focal female’s great grandoffspring). We expect stochastic effects to eventually overwhelm this intergenerational cascade, but the additional impact beyond grandoffspring has non-zero impact, as evidenced by the small differences between the deterministic results and the stochastic, agent-based results (i.e. the difference in the gold lines in Figure 2).

## Discussion

Our three related, but distinct, modeling approaches yield the same conclusion: when maternal survival has strong impacts on the survival of offspring and grandoffspring, populations evolve longer lives with less frequent reproduction.

Our models build on previous work in this area, most directly the Mother and Grandmother Hypotheses that seek to explain the adaptive value of menopause in humans and those few other species in which it has been observed (22–25). These hypotheses posit that, in species in which mothers are likely to have a large number of dependent offspring, it is to their advantage to shut down reproduction as they get older, especially as the frequency of reproduction declines and the costs of reproduction increase. These mothers benefit from focusing investment on their own survival so as to be able to guarantee that they can continue to provide resources to their offspring and grandoffspring (22–25). These hypotheses have garnered empircal support in both humans (27) and whales (9). Our models serve, then, as a more general form of the Mother Hypothesis. Presence of dependent offspring selects for longevity at the expense of reproduction in less severe forms than the complete elimination of reproduction in later life. While menopause is a highly visible phenomenon and is particularly salient to human life history, we posit that selection to extend lifespan at the expense of reproduction acts in more subtle ways and is more widespread than is menopause.

These results also build on work studying the co-evolution of intergenerational resource transfers and longevity (40, 41). When members of one generation transfer resources to the next generation (as in human societies) and the magnitude of this intergenerational transfer increases for at least some part of the adult lifespan, then there should be selection to increase survival across ages during which intergenerational transfer is non-zero. Thus, as such intergenerational transfers evolve, so too should longevity co-evolve, creating a potential feedback loop that can lead to the evolution of long-lives and extended periods of intergenerational transfer (40, 41). Our work applies this logic specifically to mothers, a salient relationship that is unviversal across mammals.

Primates live substanitally longer lives than other mammals with comparable body sizes ((14), Figure 3). Numerous non-mutually exclusive hypotheses have been put forward to explain this difference, including primates’ low rates of extrinsic mortality (22, 42), arboreality (43), large brains (44), and the unpredictability of food access (45). Our models add to this discussion by suggesting that part of the difference in lifespans between primates and non-primate mammals lifespan is explained by the strong links that exist between maternal survival and offspring fitness in many primates.

**Figure 3.**
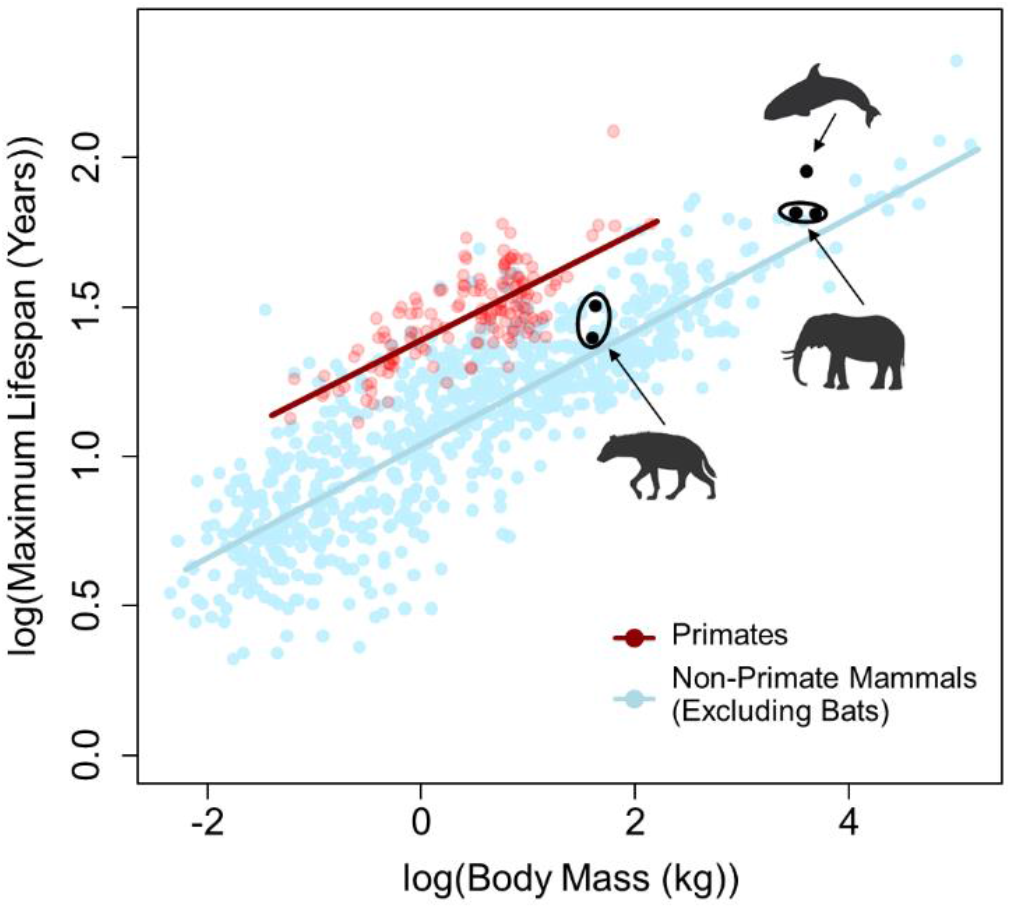
Primates live substantially longer lives than non-primate mammals of comparable body sizes (data from the AnAge database (46), excluding bats). Our models suggest that part of this difference in lifespan can be explained by links between maternal survival and offspring fitness leading to selection for longevity in mothers. Points in black indicate non-primate genera in which mother-offspring fitness links similar to those present in primates have also been documented. In each case, species of these genera live longer lives than predicted by their body sizes. Silhouettes generated via BioRender.

Although we have parameterized our models based on data from a primate species, our predictions hold for mammals, generally. We expect any species in which offspring fitness is strongly tied to maternal survival after weaning to be selected to live longer, slower lives than would otherwise be expected for their body size. Similar post-weaning mother-offspring fitness links to those parameterized in our models have been documented in several non-primate species, including spotted hyenas (10), orca whales (9), and elephants (11). In each of these three cases, species for whom maternal survival strongly influences offspring fitness live longer than would be expected based on their body size, consistent with the predictions of our models (Figure 3).

Our models may also partially explain the notable male-biased mortality that is widespread (though certainly not universal) across mammalian taxa (47, 48). Female mammals almost universally provide greater care and do so for a longer period of time than do fathers (3, 5) (though paternal care does occur in many species and can be important (49–52)). When mother-offspring fitness links exist and are stronger than the equivalent father-offspring fitness links, we predict that sexually dimorphic strategies for investing in survival and reproduction should evolve, with females assigning a relatively higher priority to survival than males. Data on father-offspring fitness links are much harder to come by than mother-offspring fitness links, but data from humans (37), baboons (53–55), and other primates (56, 57) suggest that the strength and duration of influence of mothers on offspring outcomes is stronger than that of fathers.

We close by laying out the process by which the central hypothesis we present here could be most thoroughly tested. Individual-level demographic data exist for dozens of long-term studies of wild mammals. Although there is a concentration of these studies within primates, many long-term non-primate study systems also exist (e.g. (9–11, 58–65)). For each of these study populations, researchers could readily measure the magnitude of each of the four mother-offspring fitness links in Figure 1. With these parameters in hand, along with life table data, an estimate of *λ* (Figure 2) can be readily calculated for a given *δ*_*m*_ (e.g. *δ*_*m*_ = 1.05). If increased linkage between maternal survival and offspring fitness has led to the evolution of long, slow lives, we predict that this value of *λ* should be negatively correlated with a species’ deviation from their predicted lifespan, given its body size. Thorough testing of our hypotheses is therefore readily possible via a large-scale comparative study that harnesses the demographic data that already exist from many populations.

## Acknowledgements

We gratefully acknowledge our sources of funding that made this work possible. MNZ has been supported by an NSF postdoctoral fellowship in biology (award # 2109636) and a Klarman postdoctoral research fellowship from Cornell University. Development of the ideas in this paper benefitted from conversations with Susan C Alberts and Jenny Tung. We are also grateful to all long-term researchers who have chosen to make their life table data from their population publicly accessible, as the Amboseli Baboon Research Project has done, which facilitates this sort of synergy between empiricism and theory.

## Data and Code Availability

During the review process, the code used to run the deterministic and agent-based models as well as the output data from those models that underlies Figure 2 can be accessed at the following Box folder: https://cornell.box.com/s/7awr0zfzctm6eke5xuy32vnxf5t1vu56. Prior to publication these files will be deposited in a permanent repository.

## Supplemental Material

### Complete Individual-Based Model Methods

These simulations track individual females in a population (males are ignored) and model the fixation of a mutation that influences birth and mortality rates. We evaluated two different scenarios: one without any links between maternal survival and offspring fitness (Scenario 1 in the main text) and one with all links considered in the main text (Scenario 4). Our model was parameterized to match savannah baboon populations (*Paio cynocephalus*, Bronikowski et al 2016). Our simulations had discrete time steps which approximated years in the baboon population (therefore we will refer to time steps as years from now on). Each year, the events occurred in the following order: adult mortality, population regulation, births, juvenile mortality, and then maturation (age increased by 1).

Adults (individuals older than the age of maturity) died when they reached their age of death, which was stochastically determined at birth. The age of death was determined by iteratively moving through each age in the life table after the age of maturity (juvenile mortality was determined separately, see below). For each age, a random value was drawn from a uniform distribution (0–1), if that value was less that the probability of death, that age was as assigned age of death. In Scenario 4, if an individual’s mother will die before it would reach maturity, all probabilities in the death table above the age of maturity were multiplied by *s*_*4*_ (if the new value was greater than 1, the probability was set at 1). The age of death was determined at birth because this allowed us to know whether a juvenile’s mother was going to die before it reached maturity, which was necessary for our questions.

The population was regulated so that there could only be *K* adults in the breeding population. If following adult mortality, there were more than *K* adults in the population, 80% of the population randomly died. We chose this form of population regulation so that in most years the adult population size was below *K*. All adults in the population after population regulation gave birth to one offspring with an age-specific probability given by the life table.

The baseline juvenile mortality was determined by the life table. In Scenario 4, juvenile mortality was also influenced by whether not the juvenile’s mother was alive and whether or not the juvenile’s mother’s mother was died before it reached maturity. This was implemented in the following ways: if a juvenile’s mother will die before it would otherwise reach maturity, the mortality in the life table was multiplied by *s*_*3*_; if a juvenile’s mother had experienced maternal loss before it had reached maturity, the juvenile’s mortality was multiplied by *s*_*5*_; if both the following were true, the mortality given by the life table was multiplied by *s*_3_ × *s*_5_; and if a juvenile was less than 1 year old (not yet weaned) when its mother died, the juvenile died.

To model life history evolution, we incorporated a mutation that changed both birth and death rates. For individuals carrying the mutation, the age-specific probabilities of mortality and giving birth given in by the life table were multiplied by a factor of *δ*_*m*_ and *δ*_*q*_, respectively. The simulations were initiated with *K* non-mutant individuals with randomly assigned ages. We then ran the simulation for a 200-years to allow the population to reach a stable age distribution. After this 200-year period, 50% of the individuals were changed to mutants. All offspring of non-mutants were mutants and all offspring of mutants were mutants (*i*.*e*., there were no additional mutations after mutants were added). We then ran simulations until the population was either all mutants or non-mutants or until 1,000 years of the simulation passed. We ran 1,000 replicate runs of the simulation for a range of *δ*_*m*_ and *δ*_*q*_ values and recorded the mean frequency of the mutants at the end of the simulation. For a given *δ*_*m*_, we calculated (using linear interpolation) the value of *δ*_*q*_ at which the mean frequency of the mutants at the end of the simulation was 50% (*i*.*e*., there was an equal chance of either mutants or non-mutants fixing). This value is equivalent to *λ* in the main text, and we use this value to compare our agent-based and deterministic model outcomes.

